# Host mRNA 3’-end processing machinery are critical binding partners during dengue virus infection

**DOI:** 10.1101/2025.06.20.659860

**Authors:** Min Jie Alvin Tan, Ahmad Nazri Mohamed Naim, Yan Ting Lim, Radoslaw M. Sobota, Martin L. Hibberd, Subhash G. Vasudevan

**Author notes:** address correspondence to Min Jie Alvin Tan.

## Abstract

Dengue virus (DENV) causes a wide variety of dengue diseases that threaten over half of the world’s population. At present, there is no antiviral therapeutics against dengue. Virus infection and its associated pathologies are the result of a myriad of interactions between the virus and its host, of which most are protein-protein interactions (PPIs). Virus-encoded proteins modulate the host environment for the virus’ benefit by interacting with host proteins during infection. Thus studies of these virus-host PPIs are most informative when performed in the context of an infection. We undertook a comparative affinity purification-mass spectrometry approach to generate a DENV-host protein-protein interactome during DENV infection and identified mRNA 3’-end processing protein complexes as novel interactors of DENV NS3. We demonstrated that the presence of the cleavage and polyadenylation specificity factor complex proteins, but not their enzymatic activity, is crucial for DENV replication, thus unveiling a potential new direction for the development of host-directed antiviral therapeutics.

## INTRODUCTION

Dengue virus (DENV) is estimated to infect about 400 million people worldwide every year, and over half of the world’s population (3 billion people) resides in regions that are at risk from dengue (1). It is the etiological agent of a wide variety of dengue diseases ranging from mild dengue fever to severe dengue hemorrhagic fever and potentially fatal dengue shock syndrome (2). Currently, there is no approved antiviral treatment for dengue while the available vaccines are limited in efficacy and availability.

Similar to other members of the flavivirus genus, the DENV genome consist of a single-stranded positive sense RNA of ∼11000 nucleotides encoding a ∼3400 amino acid residue polyprotein precursor that is post-translationally cleaved into three structural proteins (capsid, pre-membrane/membrane and envelope) and seven non-structural (NS) proteins (NS1, NS2A, NS2B, NS3, NS4A, NS4B and NS5) (3). Virus replication and pathogenesis are the result of the perturbations induced in the host protein network by virus proteins during virus infection. Thus the elucidation of the virus-host interactome will be instrumental to a complete understanding of the cellular processes during a virus infection, and the consequent identification of interacting host proteins that are crucial for virus replication represent potential targets for host-directed antiviral therapeutics.

Loss-of-function genetic screens have proven useful in identifying host dependency factors that are critical for DENV replication (4–7), but whether and how these factors interact with virus proteins during virus infection and pathogenesis is often not uncovered. In contrast, proteomic studies directly identify host proteins and pathways targeted by DENV proteins. Proteome-wide studies applying the expression of fragments of individual virus proteins using the yeast-2-hybrid (Y2H) method (8, 9) or individual full-length virus proteins using the affinity purification coupled with mass spectrometry (AP-MS) approach (10, 11) have had success in identifying host protein interactors. But they only reveal part of the story as these studies do not replicate cellular conditions during virus infection. In addition, it is known that the flavivirus proteins interact with one another during virus infection (12). Hence such studies are also missing out on DENV-host protein interactions that are mediated by multi DENV protein complexes.

Virus-host protein interactome studies in the context of virus infection have successfully identified critical interacting human proteins and pathways for influenza infection (13, 14). Similar studies in flaviviruses have been held back by a compact and highly sensitive flavivirus genome intolerant to manipulation (15), despite advances in the development of reverse genetics systems in the construction of stable flavivirus infectious DNA clone constructs (16, 17). Attempts at sequence insertions have often produced constructs that were less fit compared to wild type and/or unstable during continuous passaging in cells (17). This has limited studies using replicon and infectious DNA clone systems to interrogate the interactome of virus proteins to individual virus proteins (18, 19). Thus, it has remained a tremendous challenge to perform a virus proteome-wide study and comprehensively characterize the flavivirus-host protein interactome during virus infection.

In this study, we seek to overcome these challenges by co-expressing an epitope tagged DENV protein during virus infection. Following affinity purification and mass spectrometry, we identified about 200 host proteins as prospective interacting proteins (PIPs) of the DENV non-structural proteins. Proteins involved in the metabolism of RNA were highly enriched amongst these PIPs, with protein complexes involved in mRNA 3’-end processing pathway in particular being targeted by NS3. We further demonstrate that DENV requires the presence of the cleavage and polyadenylation specificity factor (CPSF) complex for replication in a manner separate from CPSF’s endonuclease activity. Collectively, our results represent a systematic DENV-human protein interactome in the context of DENV infection that has directed the discovery of an interacting host protein complex crucial for virus replication independent of its enzymatic function. This discovery opens a new angle for the development of host-targeted antiviral therapeutics.

## RESULTS

### Identification of interacting host proteins during DENV infection

We first generated expression plasmids for each virus protein using the infectious clone of a clinical strain of dengue serotype 2 virus (EDEN2) as the template (20). Flavivirus proteins are translated as a polyprotein that is subsequently cleaved by both the host proteases furin and signalase on the lumen side of the ER and the viral NS2B/NS3 protease on the cytoplasmic side to yield the individual viral proteins (Fig. 1A). Hence they do not typically start with the methionine residue, except the capsid (C) protein. Considering the possible role of the N-terminal region in the insertion of NS2A and NS4A into the membrane (21, 22), we decided against adding an N-terminal methionine residue or using a N-terminal FLAG tag to initiate translation. Instead we used the ubiquitin fusion technique to express the DENV proteins with their native N-terminal residue (23, 24). The coding sequence for a HA-tagged K48R ubiquitin (Ub) is cloned directly before the coding sequence of the flavivirus protein. Upon expression in the cell, the ubiquitin is cleaved by cellular de-ubiquitinases (DUBs), leading to the release of the DENV protein with its native N-terminal residue (Fig. 1B). Finally, we added a C-terminal FLAG tag to enable the use of a common antibody for affinity purification of all the DENV proteins. This also permits the generation of control samples that are used to normalize the background interactors during comparative analysis to identify human protein interactors specific for each DENV protein.

**Fig. 1.**
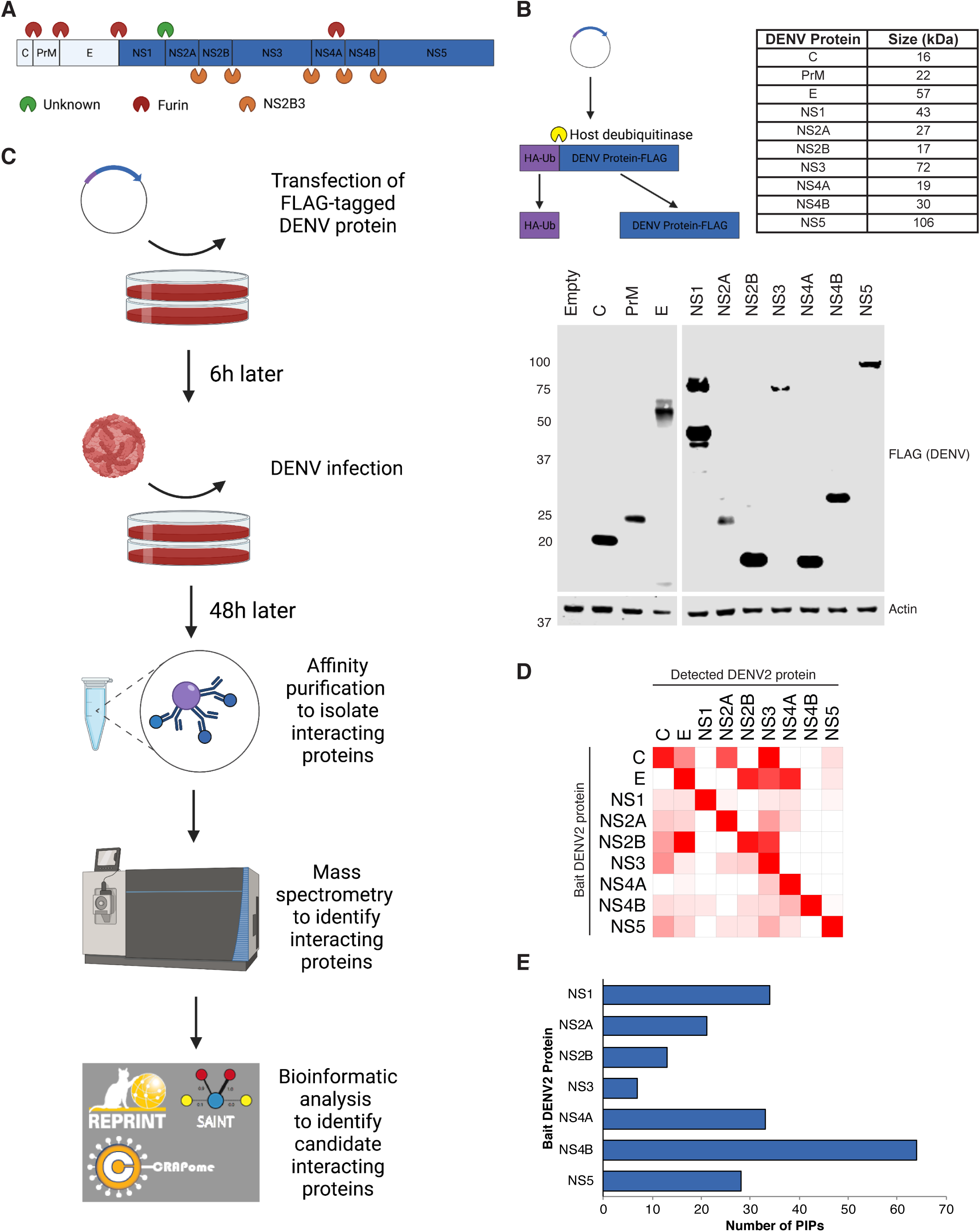
Building a virus-human protein interactome during dengue virus (DENV) infection. **a.** The DENV genome is translated as a single polyprotein that is post-translationally cleaved by host and DENV proteases to produce 10 DENV proteins. **b**. Cloning strategy for the expression of DENV proteins with their native N-terminal residue. The coding sequence for a HA-tagged K48R ubiquitin is cloned directly before the coding sequence of the flavivirus protein. Upon expression in the cell, the ubiquitin is cleaved by cellular de-ubiquitinases (DUBs), leading to the release of the flavivirus protein with the native N-terminal residue. Western blot showing the expression of the FLAG-tagged DENV proteins at their expected sizes after transient transfection in Huh7 cells (below). **c**. Schematic of the experimental setup for the co-expression of a FLAG-tagged DENV protein during DENV infection, followed by affinity purification-mass spectrometry and REPRINT analysis to score identified proteins. **d**. Heatmap depicting the average CRAPome scores of detected DENV proteins from the biological replicates of each bait DENV protein, relative to the average CRAPome score for that bait DENV protein. **e**. Bar graph showing the number of prospective interacting proteins (PIPs) identified for each DENV non-structural protein based on their SAINT scores.

We then transfected these expression plasmids into Huh7 cells to verify the expression of various DENV proteins at their respective expected sizes by Western blot (Fig. 1B). Proper and complete cleavage of the HA-Ub molecule from the DENV proteins was also confirmed (Supplementary Fig. 1). The characteristic polyubiquitin smear was observed in all the samples, except the empty vector and capsid samples, as the capsid (C) expression plasmid (second lane) was not cloned with a HA-Ub sequence since it has a native methionine.

Having established the correct expression of each virus protein, we proceeded to prepare samples for the mass spectrometry experiments. We transfected the respective plasmids individually into Huh7 cells, followed 6 hours later by infection of the cells with DENV (Fig. 1C). The FLAG-tagged bait proteins were then extracted for affinity purification 48 hours post-infection before being sent for identification by mass spectrometry together with their associated proteins. We generated triplicate affinity purification samples for each DENV bait protein but were unable to reliably detect the bait proteins in some of these replicate samples (e.g. one of the triplicates for NS2B). After omitting these replicates from our analysis, thousands of host proteins were collectively identified as being associated with the DENV bait proteins (Supplementary Table 1).

To facilitate the identification of reproducible and DENV bait protein specific interacting host proteins, we analyzed the replicate samples for each DENV bait protein using two different scoring algorithms on the Resource for Evaluation of Protein Interaction Networks (REPRINT) platform: Contaminant Repository for Affinity Purification (CRAPome) and Significance Analysis of INTeractome (SAINT) (25, 26) (Supplementary Table 1). Both algorithms compare the spectral counts of proteins detected in the respective replicate bait DENV protein samples to negative control samples generated under the same affinity purification conditions to produce an enrichment (FC for CRAPome) or probability (SP for SAINT) score.

This dataset represents a comprehensive database of the DENV-human protein interactome in the context of DENV infection. Hence we were first interested to see the number and frequency of DENV proteins for each of the bait DENV proteins (Fig. 1D). We detected extensive interactions between the DENV proteins, including previously described interactions such as those between NS3 and NS5 (27, 28), NS4A and NS4B (29), NS1 and NS4A (30, 31), and NS2A and C (32).

While this is unsurprising considering that all the non-structural proteins come together to form the replication complex (33), further studies are necessary to investigate the significance and context of these interactions. In particular, we detected a strong reciprocal interaction between the E and NS2B proteins that is likely to be occurring outside the context of the replication complex. We also noted that none of the detected DENV proteins scored as significant on either CRAPome (FC ≥ 4) or SAINT (SP ≥ 0.9) (Supplementary Table 2), suggesting that such interactions make up a relatively small proportion of the interactions that DENV proteins make during virus infection.

To increase the stringency in identifying human interacting proteins unique to each bait DENV protein, we examined the scores assigned by SAINT to the human proteins and identified about 200 proteins that were scored as significant (SP ≥ 0.9) in at least two of the three replicate samples of a bait DENV protein. We termed these proteins as prospective interacting proteins (PIPs) and found that they were well distributed amongst the various non-structural proteins (Fig 1E). In addition, more than half of the proteins that were found in all three replicates have been previously identified as an interactor or differentially expressed during DENV infection (Supplementary Table 3), validating the effectiveness of our approach in identifying true interacting proteins.

### DENV proteins interact with host proteins involved in diverse host pathways

When we visualized the list of PIPs in a DENV-human protein interaction map, we found that there were numerous human proteins that were identified as PIPs of more than one DENV protein (Fig. 2A). This finding stands out from previous proteomic studies in which there were few overlapping interacting proteins identified between the virus proteins (10, 11, 34), and could be attributed to our study being performed in the context of DENV infection that enabled the identification of host proteins binding to DENV protein complexes such as the virus replication complex.

**Fig. 2.**
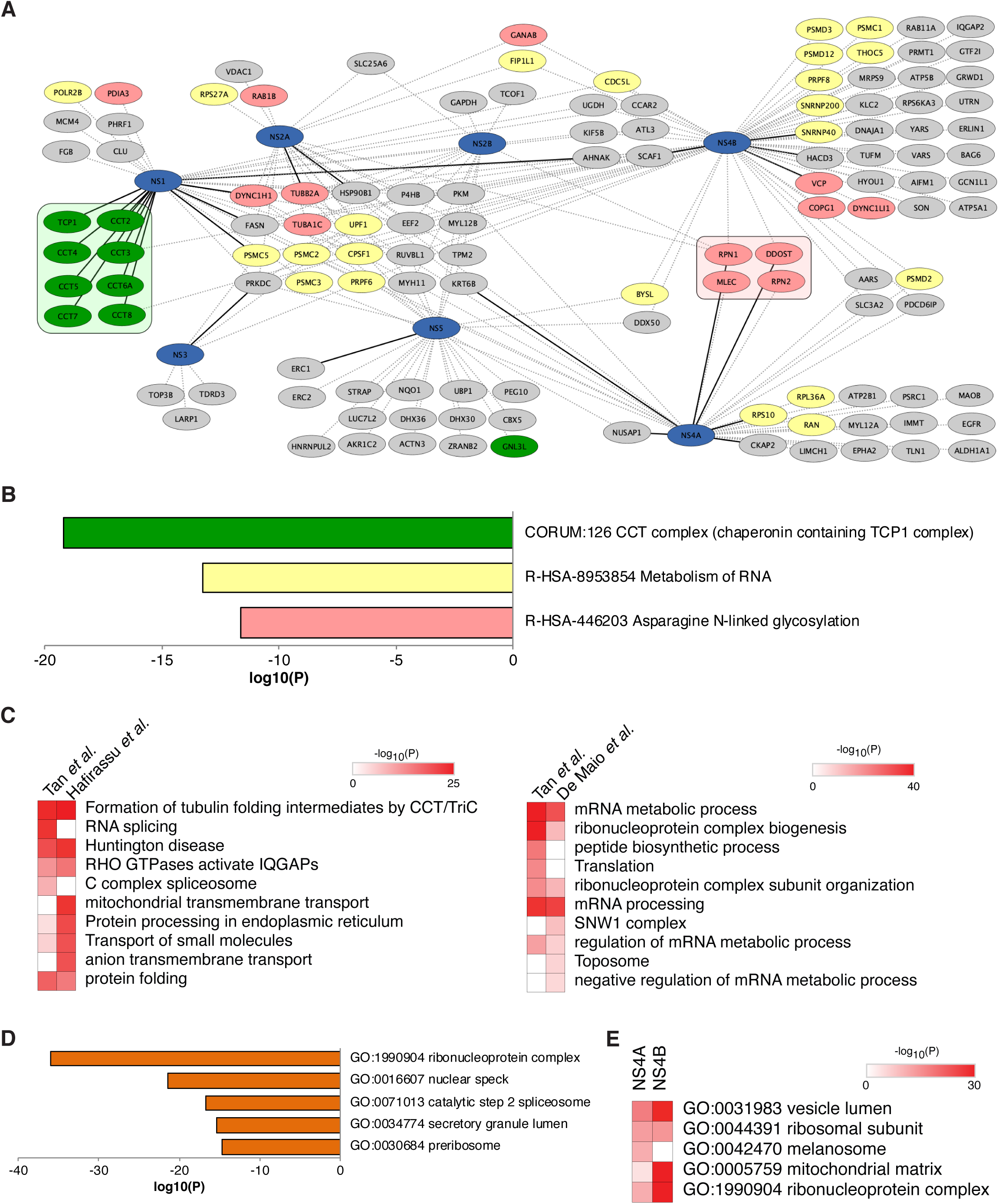
The DENV-human protein interaction map during DENV infection. **a**. The prospective interacting proteins (PIPs) (grey) of all the DENV non-structural proteins were used to construct this DENV-human protein interaction map during DENV infection. The blue nodes represent the bait DENV proteins while the green, yellow and red nodes highlight the human proteins associated with the CCT complex, RNA metabolism and asparagine N-linked glycosylation respectively. In addition, the proteins that make up the core CCT and OST protein complexes are boxed. Dotted and solid lines represent interactions scored as significant by SAINT in two or three biological replicates respectively. **b**. Bar graph showing the top three enriched pathways associated with the PIPs for all the DENV NS proteins. **c**. Heat map depicting the enriched pathways associated with human proteins pulled down by DENV NS1 (left) and NS5 (right) in this and previous studies. **d**. Bar graph showing the enriched cellular compartments associated with NS5 candidate interacting proteins using the Gene Ontology Cellular Components database. **e**. Heat map showing the enriched cellular compartments associated with NS4A and NS4B candidate interacting proteins using the Gene Ontology Cellular Components database. All the pathway enrichment data was generated with the Metascape analysis tool searching the Gene Ontology Biological Processes, CORUM, Reactome Gene Sets, KEGG Pathway and WikiPathways databases for b and c, and the Gene Ontology Cellular Components database for d and e. The magnitude of the bars (b and d) and the color intensity (c and e) represents the statistical significance of association to the respective pathways (-log_10_(P)).

We next performed pathway enrichment analysis to identify host pathways that are enriched in the list of PIPs. We found these proteins to be involved in diverse host processes (Supplementary Table 4), with the most significant enrichment in the proteins associated with the chaperonin containing tailless complex polyprotein 1 (CCT) complex (green), along with proteins involved in diverse host processes including RNA metabolism (yellow) and asparagine-linked glycosylation (red) (Fig. 2A and 2B, and Supplementary Table 4).

Two previous studies have performed AP-MS studies by incorporating epitope tags into the NS1 and NS5 gene in the context of a replicon or infectious clone (18, 19). To compare our respective approaches, we performed pathway enrichment analysis on the human proteins that were identified in both NS1 and NS5 datasets.

We found both lists of identified proteins to be significantly enriched in the same pathways (Fig. 2C and Supplementary Tables 5 and 6). In particular, the CCT complex (Formation of tubulin folding intermediates by CCT/TriC) was significantly enriched in both NS1 lists (Fig. 2C, left) while mRNA metabolic process, mRNA processing and ribonucleoprotein complex biogenesis were most significantly featured in both NS5 lists (Fig. 2C, right). Taken together, our approach compares favorably in the identification of human protein interactors with the reverse genetics approach that directly incorporates an epitope tag into the virus genome.

Finally, we checked that our exogenously expressed FLAG-tagged DENV proteins are localizing to their appropriate cellular compartments. We used the gene ontology (GO) cellular components database to analyze the list of interacting proteins for three DENV proteins (NS4A, NS4B and NS5) with previously defined subcellular localization (Supplementary Table 7). The top three terms for NS5 were ribonucleoprotein complex, nuclear specks and catalytic step 2 spliceosome (Fig. 2D), which is expected with DENV2 NS5 being an RNA-dependent RNA polymerase, its known localization to the nucleus (35) and its ability to interact with and modulate splicesome activity (19). Consistent with DENV NS4B protein’s reported localization to the mitochondria and modulation of mitochondrial structure (36, 37), the top hit from the GO cellular components analysis was the mitochondrial matrix (Fig. 2E). Finally the vesicle lumen was amongst the top hits for both membrane-bound NS4A and NS4B (Fig. 2E). The results of these analyses suggest that the expressed DENV proteins are localizing to their native cellular components within the cell and interacting with previously reported host proteins.

Collectively, we have generated a comprehensive DENV-human protein interactome in the context of DENV infection and identified host proteins and pathways targeted by DENV non-structural proteins during infection.

### Oligosaccharyltransferase (OST) complex members are a common interactor of DENV non-structural (NS) proteins

Asparagine-linked glycosylation was one of the top terms picked up by pathway enrichment analysis and is catalyzed by the oligosaccharyltransferase (OST) complex (Fig. 2B). The OST complex is an essential human host protein complex for flavivirus infection (4, 5, 38), although the nature of its interaction with the flavivirus proteins during virus infection has not been fully characterized. We detected peptides from many members of the OST complex with varying frequency and abundance in the AP-MS samples from the various DENV proteins (Supplementary Table 8). CRAPome and SAINT analysis revealed that the OST complex members were more frequently and significantly enriched in samples from the DENV non-structural proteins than the structural proteins (Fig. 3A and Supplementary Table 8). In addition, the membrane-bound NS proteins (NS1, NS2A, NS2B, NS4A and NS4B) pulled down the OST complex members with greater frequency and abundance, with NS4A interacting with the greatest number of complex members.

**Fig. 3.**
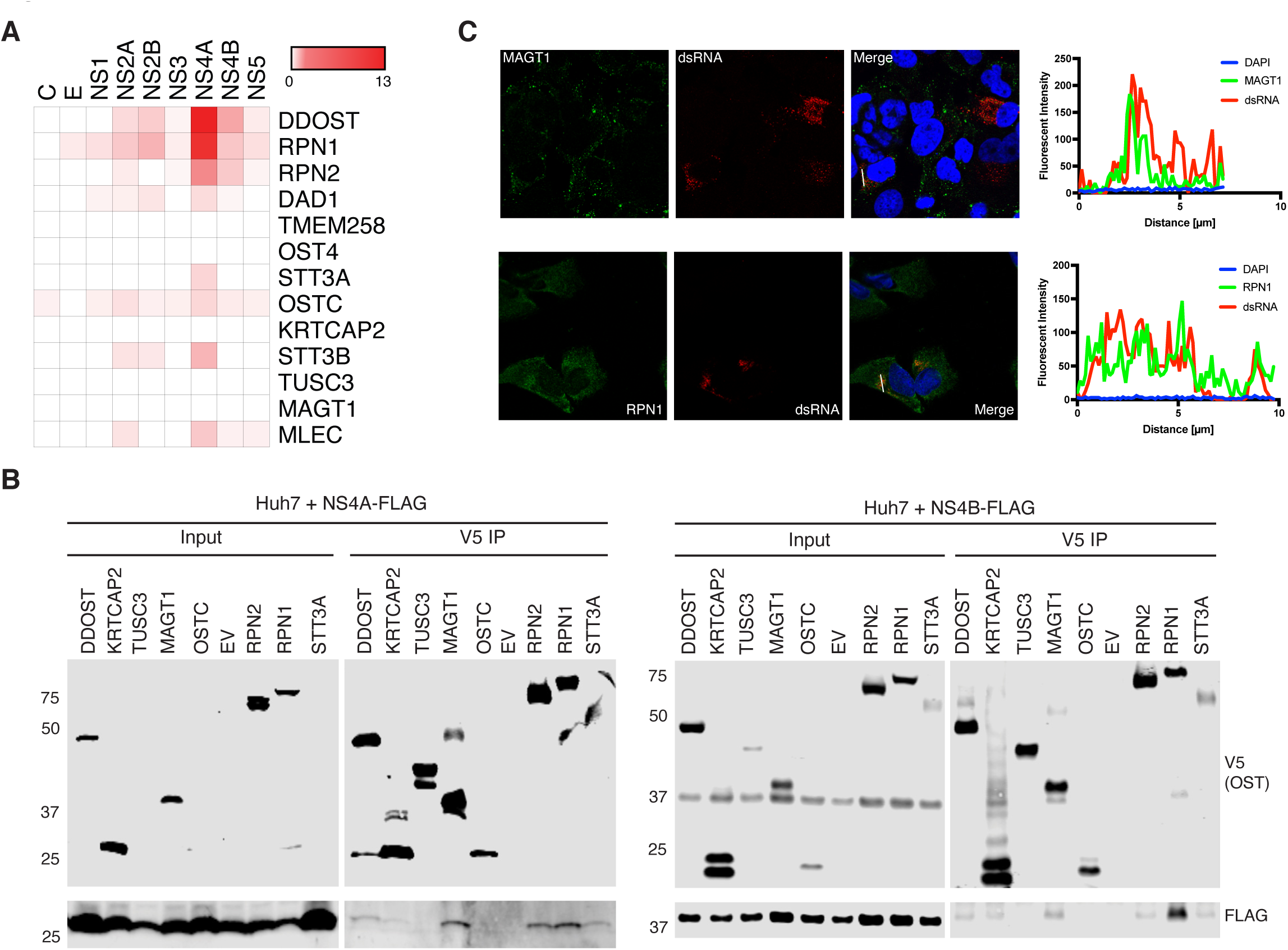
OST complex interacts with the DENV replication complex in virus replication organelles. **a**. Heat map depicting the average CRAPome scores of the detected OST complex members from the biological replicates of each bait DENV protein. **b**. V5 immunoprecipitation of 293T cells transfected with the indicated FLAG-tagged DENV proteins and V5-tagged OST complex members, followed by Western blotting with the indicated antibodies. **c**. Immunofluorescence assay of Huh7 cells infected with DENV showing the localization of dsRNA (red) and the indicated OST complex protein (green). The cells were imaged using Zeiss LSM 710 upright confocal microscope with a 63x 1.35 NA objective. The quantification and co-localization of the fluorescent intensities for the DENV and OST complex protein were performed using the Zeiss ZEN software Profile function, with the line-scan profile being graphed using Prism. The images are representative of similar results from three independent biological experiments.

We sought to validate some of these interactions by immunoprecipitation followed by Western blot (Fig. 3B). We co-transfected V5-tagged members of the OST complex and FLAG-tagged DENV2 NS4A and NS4B, and performed reverse immunoprecipitation using V5 antibodies, followed by Western blotting. Consistent with our AP-MS data, DENV2 NS4A was pulled down by DDOST, RPN1, RPN2 and STT3A (Fig. 3B, left). Interestingly, DENV NS4B was also pulled down by the same set of proteins (Fig. 3B, right).

DENV replication takes place in virus replication organelles containing double-stranded RNA and the virus replication complex (RC) made up of all the DENV NS proteins (33, 39, 40). The high frequency of OST complex members as interactors of DENV NS proteins suggests that the interaction between DENV and the OST complex could be taking place in the context of these virus replication organelles.

We examined the intracellular localization of the OST complex members RPN1 and MAGT1, and the virus RC by immunofluorescence assay (Fig. 3C). We found a strong co-localization of dsRNA with both RPN1 and MAGT1 in Huh7 cells infected with DENV, thus putting the OST complex in close proximity of the virus replication organelle. Taken together, our data supports a model in which the OST complex is recruited via interactions with DENV proteins within the DENV RC to the DENV virus replication organelle during virus infection.

### DENV NS3 interacts with and re-localizes CPSF complex proteins

RNA metabolism was the second most-enriched pathway amongst the host proteins in our DENV-human protein interactome (Fig. 2B). Amongst the DENV proteins, DENV NS3 is a multi-functional protein with helicase and nucleoside 5’-triphosphate activities. Hence, we analyzed the list of human proteins identified in our NS3 samples and found that metabolism of RNA, in particular mRNA, was the most enriched term (Fig. 4A and Supplementary Table 9). Within the list of proteins associated with this term were members of protein complexes involved in mRNA 3’-end processing. Besides the core U5 snRNP components of the spliceosome that had previously been identified as a target of NS5 (19), we also detected member proteins of the cleavage stimulation factor (CSTF), cleavage factor (CF), and the cleavage and polyadenylation specificity factor (CPSF) complexes (Fig. 4B).

**Fig. 4.**
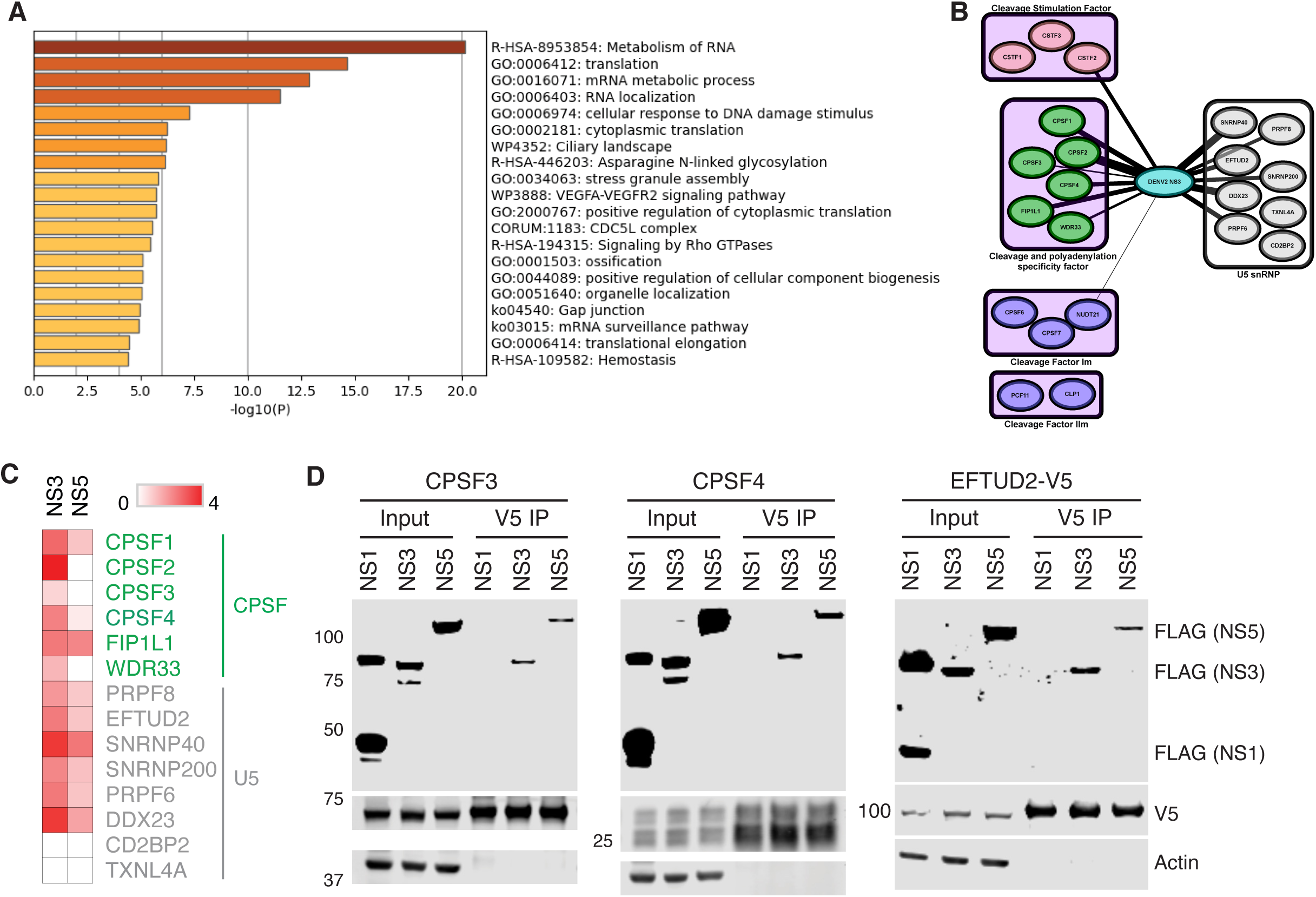
DENV targets proteins involved in multiple steps during RNA metabolism. **a**. Bar graph showing the enriched pathways associated with the human proteins identified in the DENV NS3 samples. Metascape analysis was performed as in Figure 2. **b**. Protein-protein interaction network showing the interaction between NS3 and human protein complexes involved in mRNA cleavage, polyadenylation and splicing processes. The presence of an edge denotes the detection of the human protein in the NS3 samples, and the thickness of the edge represents the average CRAPome score of the interaction from the replicate samples for NS3. **c**. Heat map depicting the average CRAPome scores of the CPSF and U5 spliceosome complex members detected in the DENV NS3 and NS5 samples. **d**. V5 immunoprecipitation of 293T cells transfected with the indicated FLAG-tagged DENV proteins and V5-tagged proteins involved in RNA metabolism, followed by Western blotting with the indicated antibodies showing the specific interaction between CPSF3, CPSF4 and EFTUD2, and DENV NS3 and NS5.

Intriguingly, some of these proteins were also detected with lower CRAPome enrichment scores in our DENV NS5 samples, suggesting that both NS3 and NS5 could be involved in interacting with members of this pathway (Fig. 4C and Supplementary Table 10).

We next sought to confirm these interactions by co-expressing DENV NS proteins with CPSF3, CPSF4 or the spliceosome protein EFTUD2 in 293T cells. After immunoprecipitation and Western blotting, we found that NS3 and NS5, but not NS1, can be pulled down by CPSF3, CPSF4 or EFTUD2 (Fig. 4D). This is in line with our mass spectrometry results, demonstrating a specific interaction between NS3 and NS5, and proteins involved in RNA metabolism.

Since all six members of the CPSF complex were detected in our NS3 samples, we chose to investigate the functional implication of this interaction. We first examined the subcellular localization of CPSF3 and CPSF4 during DENV infection using an immunofluorescence assay (IFA) (Fig. 5A). CPSF3 and CPSF4 were primarily localized in the nucleus, but we observed a decrease in the levels of CPSF proteins in DENV-infected Huh7 cells that were co-stained with DENV NS3 (Fig. 5A and 5B) or dsRNA (Extended Data 1A). Quantification of the fluorescent intensity revealed a significant decrease in the CPSF protein levels in the nucleus, and a small but significant increase in CPSF protein levels in the cytoplasm (Fig. 5B). To investigate whether DENV NS3 alone is sufficient to re-localize the CPSF proteins, we expressed DENV NS3 in Huh7 cells and performed the same IFA. We found that Huh7 cells expressing NS3 also exhibited a significant re-localization of the CPSF proteins from the nucleus to the cytoplasm (Fig 5C and 5D). In contrast, DENV NS1 did not have a significant effect in the subcellular localization of the CPSF proteins (Extended Data Fig 1B).

**Fig. 5.**
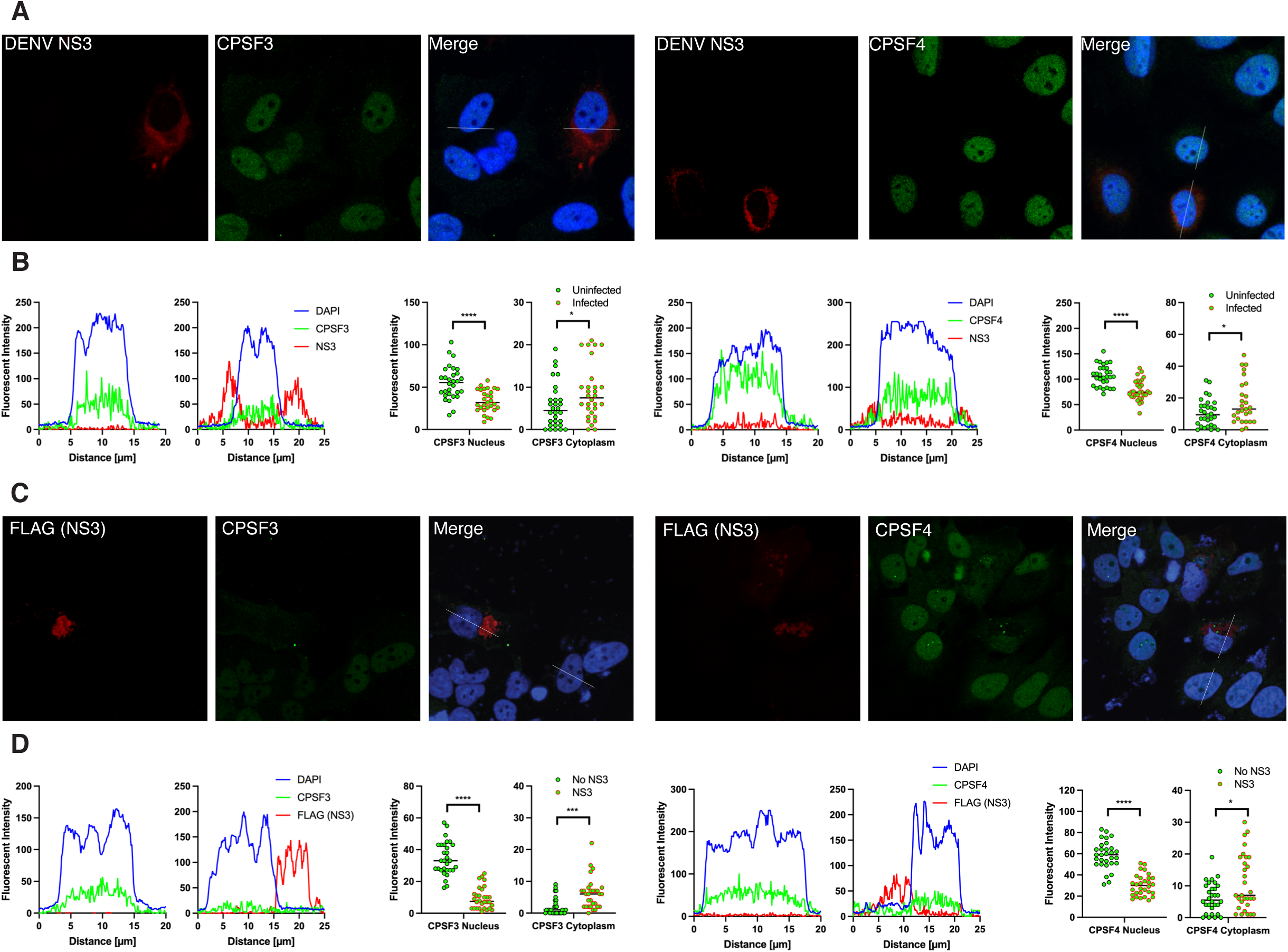
DENV NS3 relocalizes CPSF proteins from the nucleus to the cytoplasm. **a.** Immunofluorescence assay of Huh7 cells infected with DENV showing intracellular distribution of CPSF3 or CPSF4 (green) with infected cells indicated by staining with NS3 (red). **b.** Quantification of the fluorescent intensity for CPSF proteins (green), DENV NS3 (red) and DAPI (blue) along the indicated line-scans was performed using the Zeiss ZEN software Profile function and plotted on a line graph using Prism. **c.** Immunofluorescence assay of Huh7 cells expressing DENV NS3 showing the intracellular distribution of CPSF3 or CPSF4 (green) with cells expressing NS3 indicated by straining with NS3’s FLAG tag (red). **d.** Quantification of the fluorescent intensity for CPSF proteins (green), FLAG (red) and DAPI (blue) along the indicated line-scans was performed using the Zeiss ZEN software Profile function and plotted on a line graph using Prism. For b and d, thirty individual data points of the fluorescent intensity for the CPSF proteins were taken from the middle of the nucleus and cytoplasm, and compared by multiple unpaired t-tests and a p-value less than 0.05 was considered significant (*, p < 0.05; ***, p < 0.001; ****, p < 0.0001). The images are representative of similar results from three independent biological experiments.

### The CPSF complex, but not its endonuclease activity, is important of flavivirus replication

CPSF3 is the endonuclease responsible for the CPSF complex’s function of cleaving pre-mRNA before polyadenylation during the maturation of host mRNA (41). It has also been shown to be a promising drug target in some cancers and parasites (42). Influenza virus and herpes simplex virus disrupt CPSF complex functions via interactions with their respective encoded virus proteins (43). Hence we examined whether DENV NS3 and NS5 have an effect on the pre-mRNA 3’-end processing (44). We found that expression of DENV NS3 and/or NS5 did not have an effect of IL8 pre-mRNA processing. In comparison, treatment of cells with the CPSF3 enzymatic inhibitor JTE-607 led to a significant accumulation of IL8 pre-mRNA (Extended Data Fig. 2).

We next investigated whether the CPSF complex plays a role in DENV replication. We used RNA interference (RNAi) to knock down CPSF3 expression levels and then infected the cells to interrogate the importance of the CPSF complex in DENV replication. Using two unique small interfering RNAs (siRNAs) against CPSF3, we found a significant reduction in virus RNA replication (Fig. 6A). In addition, this reduction is dose-dependent, as increasing amounts of transfected siRNA that corresponded with reductions in the levels of CPSF3 RNA and protein led to decreasing amounts of virus RNA replication and virus production (Fig. 6B).

**Fig. 6.**
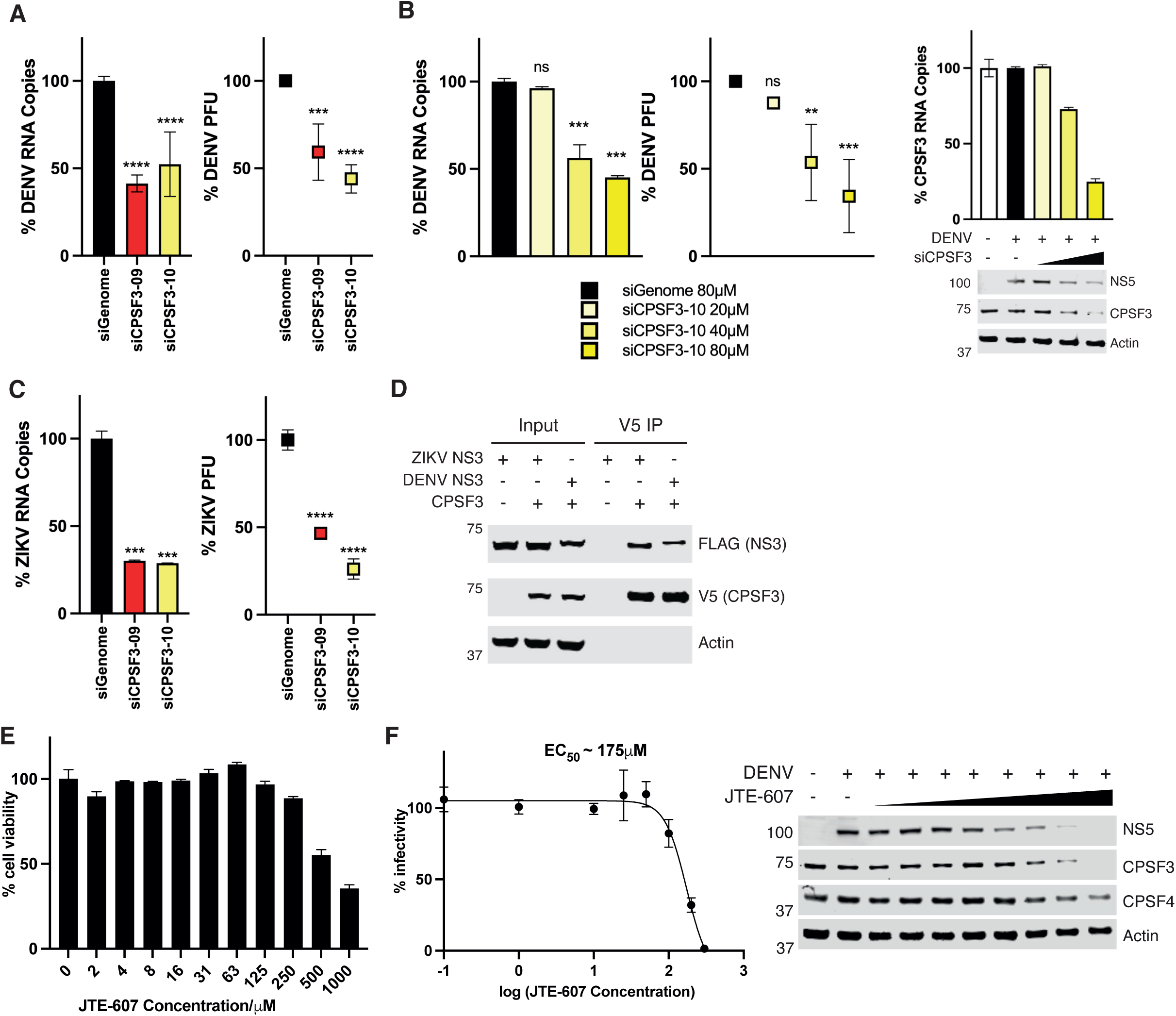
The CPSF complex, but not its endonuclease activity, is critical for DENV replication. **a**. Bar graph (left) and dot plot (right) showing the relative levels of DENV genomic RNA and DENV plaque forming units (PFU) in Huh7 cells treated with the indicated siRNAs at 40μM for 24h before being infected with DENV for 24h. **b.** Bar graph (left) and dot plot (center) showing the relative levels of DENV genomic RNA and DENV plaque forming units (PFU) in Huh7 cells treated with increasing concentrations of siCPSF3-10. Bar graph showing the relative levels of CPSF3 mRNA and Western blot with the indicated antibodies showing the protein levels of DENV NS5 and CPSF3 (right). **c**. Bar graph (left) and dot plot (right) showing the relative levels of ZIKV genomic RNA and ZIKV plaque forming units (PFU) in Huh7 cells treated with the indicated siRNAs at 40μM for 24h before being infected with ZIKV for 24h. **d**. V5 immunoprecipitation of 293T cells transfected with FLAG-tagged ZIKV or DENV NS3 and V5-tagged CPSF3, followed by Western blotting with the indicated antibodies, showing that ZIKV NS3, like DENV NS3, interacts with CPSF3. **e.** Bar graph showing the relative cell viability of Huh7 cells treated with the indicated concentrations of JTE-607. **f**. Line graph showing the relative DENV infectivity of Huh7 cells treated with the indicated concentrations of JTE-607. The number of plaque forming units were counted for each sample and plotted relative to the untreated sample. The estimated EC_50_ was calculated using Prism. Western blot with the indicated antibodies of the corresponding cell lysate samples showing the decrease in protein levels of DENV NS5, CPSF3, CPSF4 and actin with treatment of increasing concentrations of JTE-607. Mean values were compared to the respective control samples by Dunnett multiple comparison after one-way ANOVA, and a p-value less than 0.05 was considered significant (*, p < 0.05; **, p < 0.01; ****, p < 0.0001).

The OST and CCT complexes have been previously found to be crucial pan-flavivirus host factors. Hence we examined whether the CPSF complex is also important for ZIKV infection. When we knocked down expression of CPSF3 with the same CPSF3 siRNAs, we found that ZIKV replication is also abrogated (Fig. 6C). We then investigated whether the CPSF complex is targeted in a similar manner by ZIKV by testing whether ZIKV NS3 also interacts with CPSF3. We co-transfected V5-tagged CPSF3 with FLAG-tagged DENV and ZIKV NS3, and subjected the cell lysates to immunoprecipitation and Western blotting. We found that ZIKV NS3, like DENV NS3, is also able to interact with CPSF3, suggesting a common mode of targeting the CPSF complex amongst the flaviviruses (Fig. 6D).

JTE-607 is a small molecule inhibitor of the CPSF complex that binds to CPSF3 and blocks its endonuclease activity at 10μM (44, 45). We found that JTE-607 has very low toxicity with little effect on cell viability up to 250μM (Fig. 6E). We next tested the effect of JTE-607 on DENV replication and found that JTE-607 had no effect on DENV replication at an inhibitory concentration of 10μM (Fig. 6F, left). But we found that JTE-607 becomes effective in inhibiting DENV replication as we increased the concentration up to 250μM in our dose-response study, with an estimated 50% effective concentration (EC50) of ∼175μM (Fig. 6F). When we sought to understand the cause of this inhibition, we found a dose-dependent decrease in CPSF3 levels with increasing concentration of JTE-607 treatment. Interestingly, CPSF4 levels also decreased in a similar manner (Fig. 6F), suggesting that a decrease in CPSF complex levels is responsible for the observed inhibition.

Taken together, DENV targets the CPSF complex during DENV infection and re-localized CPSF proteins to the cytoplasm. The enzymatic function of the CPSF complex is not targeted by DENV and does not play a role in this abrogation, as inhibition of CPSF3’s endonuclease activity with a small molecule inhibitor has no effect on DENV replication. But a reduction in CPSF complex protein levels, whether by RNA interference or chemical treatment, leads to an abrogation of DENV replication

## DISCUSSION

During virus infection, virus proteins disrupt host protein networks to promote virus replication and pathogenesis. Hence the revelation of the virus-host interactome is a critical piece in uncovering the cellular processes that occur during a virus infection. Previous proteomic studies on DENV have been limited to studying a single DENV protein in the context of DENV infection or the study of the DENV proteins in the absence of DENV infection. Here we co-expressed epitope-tagged DENV proteins during DENV infection to generate a comprehensive DENV-human protein interactome in the context of DENV infection using an affinity purification-mass spectrometry approach (Fig. 1C). We identified about 200 human proteins as prospective interacting proteins (PIPs) of the non-structural (NS) proteins of DENV. Pathway enrichment analysis of these proteins reveals an enrichment of members of the chaperonin containing tailless complex polyprotein 1 (CCT) complex, and proteins involved in asparagine N-linked glycosylation and RNA metabolism (Fig. 2B).

The primary protein complexes responsible for the former two pathways have have been previously reported to be crucial host factors for DENV replication. The oligosaccharyltransferase (OST) complex catalyzes the co-translational glycosylation of specific asparagine residues, and had been previously identified to be an indispensable human host protein complex for flavivirus infection via genetic and genomic studies (4, 5, 38). But the nature of its interaction with the virus at a protein level had been incomplete and seemingly conflicting. We clarify this by detecting members of the OST complex in our AP-MS samples for every single non-structural (NS) DENV protein (Fig. 3A) and showing co-localization of OST complex members with dsRNA in DENV-infected cells (Fig. 3C). Taken together, these data suggest that the interaction with the OST complex is taking place in the context of the virus replication complex within the virus replication organelles. In contrast, the CCT/TRiC complex is only targeted by the NS1 protein as members of the CCT complex were only scored as PIPs for NS1. This was confirmed via immunoprecipitation followed by Western blotting, as NS1, but not NS3 or NS5, is able to pull down CCT2 and CCT7 (Extended Data Fig. 3 and Supplementary Table 11).

The RNA metabolism pathway comprises all the steps in which RNA transcription products are further modified to yield mature mRNA, including capping, splicing, 3’-cleavage and polyadenylation. We found members of protein complexes involved in splicing and mRNA 3’-end cleavage in the AP-MS samples for NS3 and NS5. In particular, we identified and validated members of the CPSF complex as novel interactors of NS3 (Fig. 4). DENV infection and the expression of DENV NS3 alone is sufficient to re-localize CPSF proteins from the nucleus, suggesting a recruitment of CPSF proteins to the cytoplasm where virus replication occurs (Fig. 5). When CPSF3 expression is knocked down using RNA interference, DENV replication was abrogated in a dose-dependent manner, indicating the importance of the CPSF complex during virus replication (Fig. 6B). The endonuclease function of CPSF3 is not involved in this role, as a small molecule inhibitor JTE-607 has no effect on DENV replication at enzyme-inhibitory concentrations (Fig. 6E). Intriguingly, we found that higher concentrations of JTE-607 led to an abrogation of DENV replication that corresponded to a reduction in the protein levels of not only CPSF3 but also CPSF4. Taken together, our results demonstrate that the presence of the CPSF complex proteins is critical for DENV replication and support a model in which DENV is targeting and recruiting the CPSF complex, but not its enzymatic activity, during virus replication.

The identification of host targets for antiviral therapeutics have been driven by genetic and small molecule inhibitor screens that have largely limited discovery to non-essential and enzymatic proteins respectively. Using a proteomic approach, we have identified a host protein complex that is crucial for DENV infection independent of its enzymatic activity. This raises the possibility of targeting essential enzymatic proteins for antiviral therapeutics by disrupting the interaction between the virus and human protein without affecting the host protein’s enzymatic activity. The potential of such an approach had been recently demonstrated in an inadvertent manner in the discovery of a dengue virus inhibitor that was subsequently found to target the NS3-NS4B interaction (46). We have found that the helicase domain of NS3 is sufficient to mediate the interaction with an N-terminal fragment of CPSF3 (Extended Data Figure 4). This finding lays the foundation for further structure-function studies necessary to uncover the molecular mechanism of the interaction, and to direct the development of specific small molecules to disrupt this interaction.

## ACKNOWLEDGEMENTS

We thank members of the Vasudevan and Sobota laboratories for discussion, advice and reagents. This work was supported by National Medical Research Council grants NMRC/OFYIRG/0006/2016 to M.J.A.T. and NMRC/CBRG/0103/2016 to S.G.V., and Ministry of Health grants MOH-000086 (MOH-OFIRG18may-0006/2019) and MOH-001497 (MOH-OFIRG23jul-0004) to S.G.V.

## AUTHOR CONTRIBUTIONS

M.J.A.T. and A.N.M.N. performed the cloning and affinity purification of the dengue virus proteins. Y.T.L and R.M.S. performed mass spectrometry and initial data analysis. M.J.A.T. performed the subsequent Metascape analysis and downstream biological assays. M.J.A.T., S.G.V. and M.L.H. supervised the research. M.J.A.T. wrote the manuscript with input from S.G.V and M.L.H.

## METHODS

### Cell lines and viruses

uh-7 and 293T (ATCC) were maintained in high-glucose Dulbecco’s modified Eagle’s medium (DMEM) containing L-glutamine supplemented with 10% [v/v] fetal bovine serum (FBS) and 1% [v/v] penicillin and streptomycin (P/S). BHK-21 cells (ATCC) were maintained in RPMI1640 medium supplemented with 10% FBS and 1% P/S. All cells were grown at 37°C in 5% CO_2_. C6/36 cells (ATCC) were maintained in RPMI1640 medium supplemented with 25 mM HEPES, 10% FBS and 1% P/S at 28°C, in the absence of CO_2_.

The two ZIKV strains, H/PF/2013 (KJ776791.2) and Paraiba01/2015 (KX280026.1), used in this study were grown in C6/36 cells, titered in BHK-21 cells and stored at -80°C (47)

The DENV strain (GenBank accessions: DENV2, EU081177) was obtained from the Early Dengue infection and outcome (EDEN) study (48). The ZIKV strain H/PF/2013 (GenBank accession: KJ776791.2) was a gift from Cécile Baronti at Aix Marseille Université (47). The viruses used in this study were grown in C6/36 cells, titered in BHK-21 cells and stored at -80°C. Institutional approval has been granted by Duke-NUS Medical School to perform experiments with DENV and ZIKV.

### Plasmid construction

The virus protein expression plasmids were generated from the DENV2 infectious clone plasmid (35) for the DENV strain. The list of primers used for plasmid cloning are listed in Supplementary Table 12. The expression plasmids for the human genes were obtained from the CCSB-Broad Lentiviral Expression Library (PerkinElmer) (49).

### Plasmid transfection and virus infection

For immunoprecipitation assays, 1.5e5 Huh7 cells were seeded onto 12-well plates and incubated overnight at 37°C with 5% CO_2_. Cells were transfected with 1μg of the indicated plasmids using FuGENE 6 (Promega), according to the manufacturer’s protocol.

For immunofluorescence assays, 1.5e5 Huh7 cells were seeded onto 12-well plates containing #1.0 glass cover slips, and incubated overnight at 37°C with 5% CO_2_. Cells were transfected with 1μg of the indicated plasmid using FuGENE 6 (Promega), according to the manufacturer’s protocol. For virus infection, cells were incubated with 500μl of virus inoculum at the indicated MOI for 1h, and viral inoculums were replaced with growth media prior to 24h incubation at 37°C with 5% CO_2_.

### Immunofluorescence assay and confocal laser scanning microscopy

The cells on coverslips were fixed with 4% paraformaldehyde for 15min at room temperature, washed with 1x phosphate-buffered saline (PBS) buffer followed by blocking and permeabilization using 2% [w/v] bovine serum albumin (BSA)/0.3% [v/v] Triton X-100/PBS for 1h at room temperature. Cells were washed with 1x PBS before incubation with the following primary antibodies: dsRNA (1001050; Exalpha), DENV NS3 (3F8) (50), CPSF3 (H00051692-M01; Novus Biologicals), CPSF4 (15023-1-AP; Proteintech), RPN1 (198598; Abcam) and MAGT1 (17430-1-AP; Proteintech). After washing with 1x PBS, the cells were incubated with the following secondary antibody coupled to Alexa-Fluor 488 and 594 (Thermo Fisher). Cell nuclei were visualized using DAPI (D9542; Merck). Coverslips were mounted using ProLong Gold Antifade reagent (Thermo Fisher) onto glass slides and confocal laser scanning microscopy was performed using a Zeiss LSM 710 upright confocal microscope (Carl Zeiss, Germany) with a 63x 1.35 NA objective. Image processing was performed with Zeiss ZEN and ImageJ software (51).

### Affinity purification-mass spectrometry sample preparation

2.5e6 Huh7 cells were transfected with 10µg of empty vector or FLAG-tagged DENV proteins. 6h post-transfection, the cells were infected with DENV as above. 48h post-infection, cells were washed twice with PBS, and lysed with 0.5% NP-40 lysis buffer (50mM Tris-HCl pH 7.8, 150mM NaCl) or 0.5% DDM lysis buffer (20mM Tris-HCl pH 7.5, 100mM NaCl). Affinity purification was performed using anti-FLAG antibody (F1804; Merck) and SureBeads Protein A/G Magnetic Beads (Bio-Rad)

### Mass spectrometry

Bound proteins were reduced with 20mM TCEP for 20min at 55°C and alkylated with 55 mM 2-chloroacetamide (CAA) for 30 min, in the dark, at room temperature. Before digestion, samples were diluted with 100 mM triethylamonium bicarbonate buffer (TEAB). Protease digestion was carried out with LysC enzyme (Wako) for 4 h, followed by trypsin (Promega) treatment for 18 h at 25°C (1:100, enzyme:protein ratio). Subsequently, samples were acidified with 1% trifluoroacetic acid and peptides were desalted with Sep-Pak C18 cartridges (Waters). Elution of peptides was performed with 0.5% acetic acid, 80% acetonitrile followed by peptide concentration using a vacuum concentrator system (Eppendorf).

### Mass spectrometry acquisition and data processing

Digested peptide samples were loaded on a heated (50°C) Easy-Spray 75µm x 50cm column (Thermo Scientific) on an EASY-nLC 1000 system coupled to a Fusion Mass Spectrometer (Thermo Scientific) with an EASY-Spray source. Peptides were separated over a 100 min gradient, using mobile phase A (0.1% formic acid in water) and mobile phase B (0.1% formic acid in 95% acetonitrile). Mass spectra were collected in data-dependent mode with cycle time of 3s between master scans. MS1 scans were performed in the Orbitrap with 120K resolution, AGC target of 400,000 and maximum injection time of 100ms. MS/MS collision induced dissociation was performed in the ion trap, with collision energy at 35 %, isolation window of 1.0 m/z, AGC target of 8,000, and maximum injection time of 70ms.

Mass spectra raw files were searched with Mascot in Proteome Discoverer 2.4 against a human uniprot database (retrieved in 2016), with following parameters: precursor mass tolerance of 10ppm, fragment mass tolerance of 0.8 Da, trypsin as enzyme with maximum 3 missed cleavages, with static modification for carbamidomethyl (C), and dynamic modifications for acetyl (Protein N-terminus), oxidation (M) and deamidation (N,Q). False discovery rate estimation with 2 levels: Strict = FDR 1%, Medium = FDR 5%. Precursor mass peak (MS1) intensities were quantified by label-free quantification (LFQ) using the Minora feature detector.

### Data availability

The raw spectra and search data will be uploaded to the Japan ProteOme STandard Repository (jPOSTrepo).

### Interactome analysis

The lists of detected proteins for each DENV bait protein was analyzed on the Resource for Evaluation of Protein Interaction Networks (REPRINT) platform using the Significance Analysis of Interactome (SAINT) and Contaminant Repository for Affinity Purification (CRAPome) analysis tools (25, 26), and prey proteins with a SAINT score ≥ 0.9 in more than two of the three replicates for its DENV bait protein were designated as prospective interacting proteins (PIPs). The DENV-host protein interactome was visualized using Cytoscape (52). The complete details of the analyses are provided in the supplementary data (Supplementary Table S6).

### Enrichment analysis

Each list of candidate interacting proteins was analyzed with a number of ontology sources including KEGG pathway, gene ontology biological processes, CORUM, reactome gene sets and gene ontology cellular components. All the pathway enrichment analysis and visualizations were performed using Metascape (http://metascape.org/gp/index.html#/main/step1) (53). The complete lists containing the terms, results and statistics for all the Metascape analyses are provided in Supplementary Information.

### RNA extraction and RT-qPCR

Total RNA from cell lysates was isolated using the RNeasy kit (Qiagen) and subjected to cDNA synthesis using the High-Capacity cDNA Reverse Transcription Kit (Thermo Fisher) according to manufacturer’s instructions. RT-qPCR was conducted using the iQ SYBR Green Supermix (Bio-Rad) for detection of the indicated genes. Gene expression was normalized to RPLP1 expression and presented as a percentage relative to empty vector control (for the pre-mRNA assay) or untreated control (for DENV and ZIKV infection). The primers used for RT-qPCR are listed in Supplementary Table 12.

### Western blotting

Proteins were separated using NuPAGE Bis-Tris gels (Thermo Fisher Scientific) and transferred onto PVDF membranes (Merck). After blocking in 4% [w/v] BSA/0.05% [v/v] Tween-20/PBS, blots were incubated with the following primary antibodies diluted in 2% [w/v] BSA/0.05% [v/v] Tween-20/PBS: actin (MAB1501; Merck), V5 (13202; Cell Signaling), FLAG (F1804; Merck), HA (ROAHAHA; Merck), CPSF3 (H00051692-M01; Novus Biologicals), CPSF4 (15023-1-AP; Proteintech) and NS5 (GTX133327; GeneTex). Bound primary antibodies were incubated with AffiniPure donkey anti-rat, mouse or rabbit Alexa Fluor 680 or 790 secondary antibodies (Jackson ImmunoResearch), followed by visualization using the Odyssey CLx Imaging System (Li-Cor).

### Cell viability and virus inhibition assays

2×10^4^ Huh7 cells were seeded onto a 96-well white opaque plate (Grenier) and treated with JTE-607 at the indicated concentrations for 24h. Cytotoxicity was determined using the CellTiter-Glo Luminescent Cell Viability Assay (Promega) according to manufacturer’s instructions. The cell viability curve is presented as a percentage of the luminescence units detected from the treated sample to that of the untreated control sample. 1×10^5^ Huh7 cells were infected with DENV at MOI 1 (multiplicity of infection) for 1h, followed by treatment with the compounds at the indicated concentrations for 24h. Supernatants were collected, clarified and subjected to plaque quantification. The efficacy of JTE-607 (EC50: concentration at which the virus infection is reduced by 50%) was determined by the sigmoidal dose response curve of virus titer against drug concentration using Prism (GraphPad).

### siRNA experiments

1×10^5^ Huh7 cells were transfected with the indicated concentration of siRNA using DharmaFECT 1 (Horizon) for 24h before infection with DENV at MOI 1 for an additional 24h. Supernatants were collected, clarified and subjected to plaque quantification 48h post transfection, while RNA was extracted from the cells for RT-qPCR analysis. The ON-TARGETplus CPSF3 siRNA (J-006365-09-0002 and J-006365-10-0002) were purchased from Horizon.

### Plaque assay

2×10^5^ BHK-21 cells were seeded onto a 24-well plate and incubated overnight at 37°C with 5% CO_2_. Virus samples were diluted with RPMI 1640 serum-free media and the cells were incubated with the diluted inoculum for 1h. Virus inoculum was then removed, and the cells were overlaid with 0.8% carboxylmethyl cellulose (CMC) and further incubated for 4 (for ZIKV) or 5 (for DENV) days. The cells were then fixed with 3.7% formaldehyde and stained with 1% crystal violet for plaque visualization.

### Statistical analysis

The error bars in the figures represent the standard error of mean for each set of data calculated using Prism (GraphPad). The statistical significance of the differences between the various experimental conditions and the control was evaluated by the indicated statistical tests using Prism (GraphPad). Data from at least two biological replicate experiments were used in the statistical tests, and P values ≤ 0.05 were considered to be statistically significant.

## FIGURE LEGENDS

**Extended Data Fig. 1.**
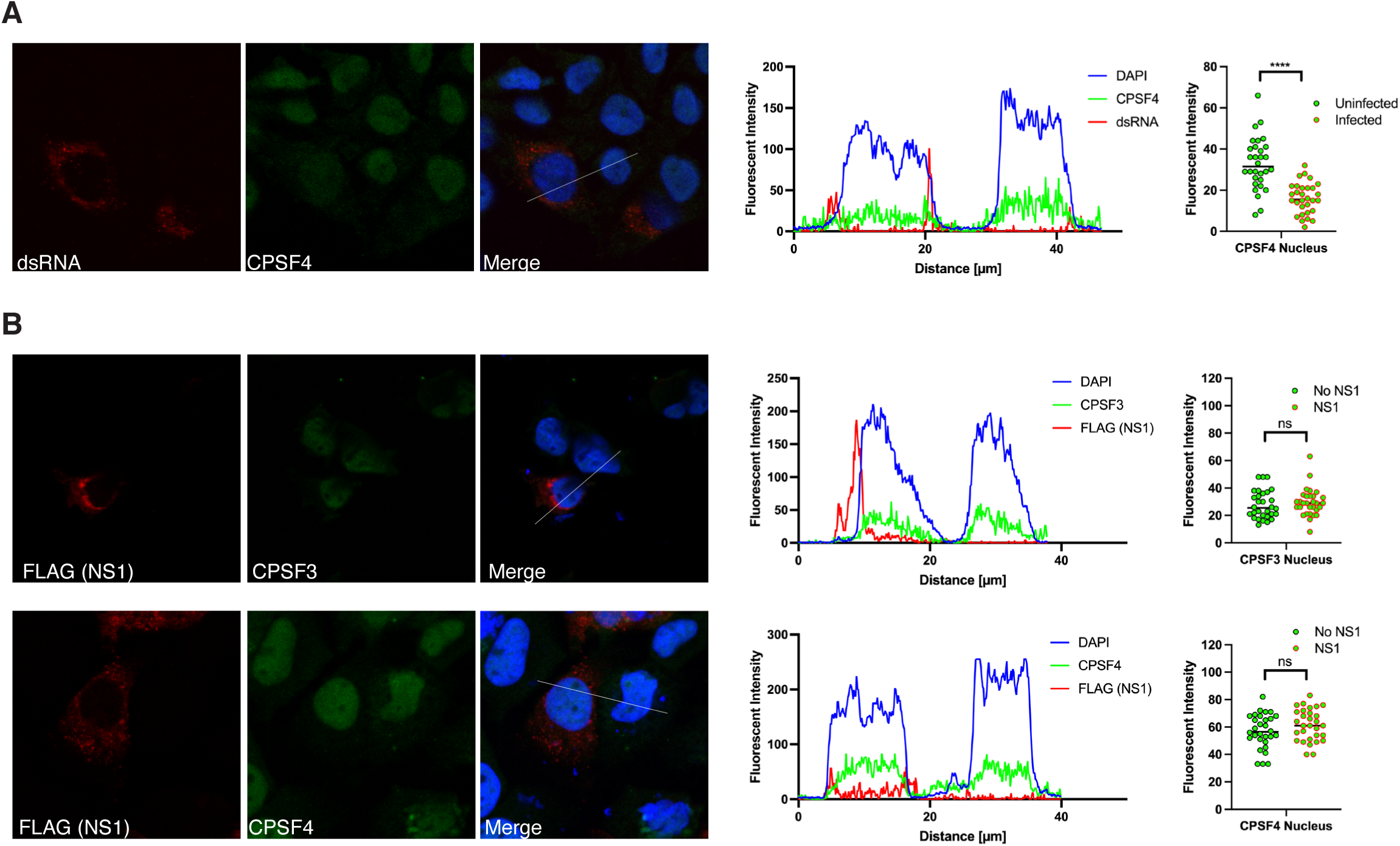
DENV NS1 does not alter the subcellular localization of CPSF proteins. **a**. Immunofluorescence assay of DENV-infected Huh7 cells showing the intracellular distribution of CPSF4 (green) with infected cells indicated by staining with dsRNA (red). Quantification of the fluorescent intensity for CPSF proteins (green), dsRNA (red) and DAPI (blue) along the indicated line-scans was performed using the Zeiss ZEN software Profile function and plotted on a line graph using Prism (right). **b.** Immunofluorescence assay of Huh7 cells expressing DENV NS1 showing the levels of CPSF4 (green) in the nuclei of cells stained for the FLAG tag on NS1 (red). Quantification of the fluorescent intensity for CPSF proteins (green), FLAG (red) and DAPI (blue) along the indicated line-scans was performed using the Zeiss ZEN software Profile function and plotted on a line graph using Prism (right). Thirty individual data points of the fluorescent intensity for the CPSF proteins were taken from the middle of the nucleus and cytoplasm, and compared by multiple unpaired t-tests and a p-value less than 0.05 was considered significant (****, p < 0.0001). The images are representative of similar results from three independent biological experiments.

**Extended Data Fig. 2.**
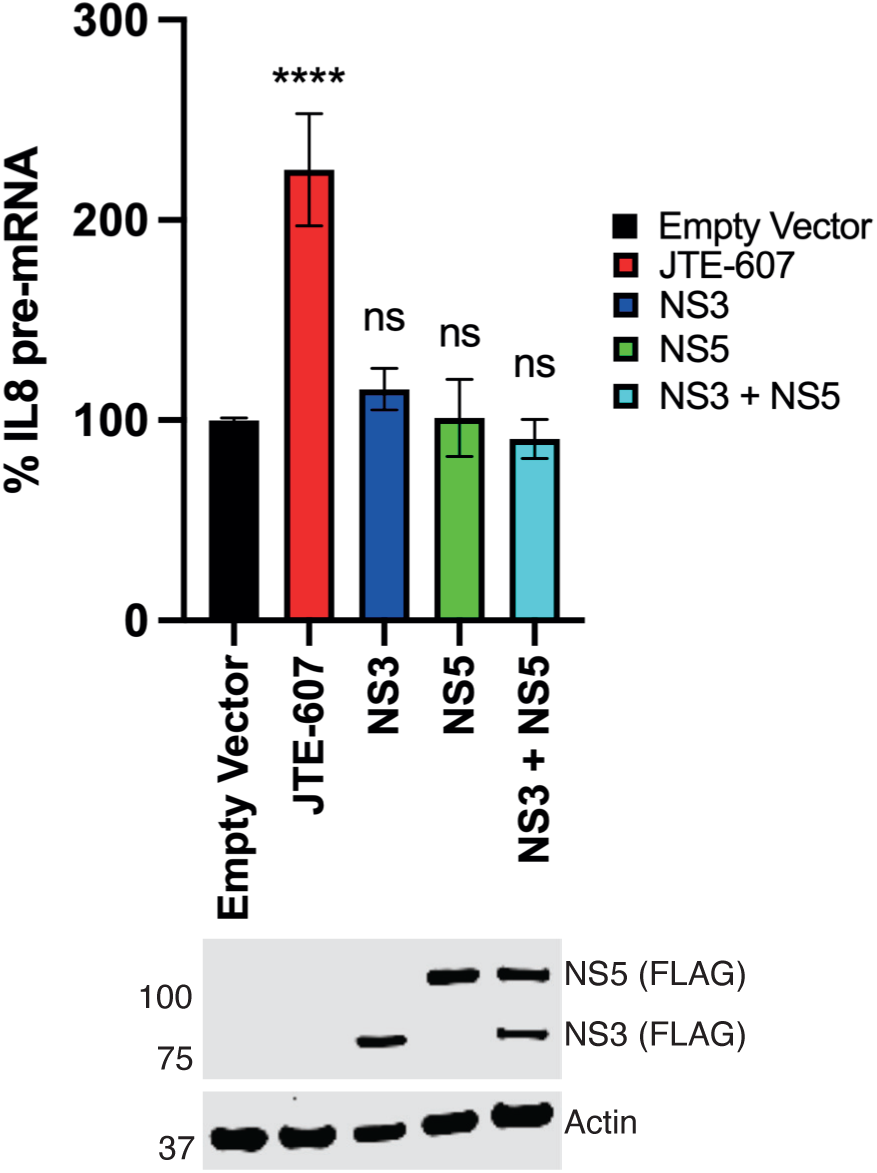
DENV NS3 does not affect CPSF complex function. Bar graph showing the relative levels of IL8 pre-mRNA in Huh7 cells treated and transfected as indicated (top). Results are from two biologically independent experiments with the statistical significance calculated by comparing mean values to the control samples by Dunnett multiple comparison after one-way ANOVA. A p-value less than 0.05 was considered significant (****, p < 0.0001). Western blot with the indicated antibodies showing the expression of NS3 and/or NS5 (bottom) is representative of two biologically independent experiments repeated with similar results.

**Extended Data Fig. 3.**
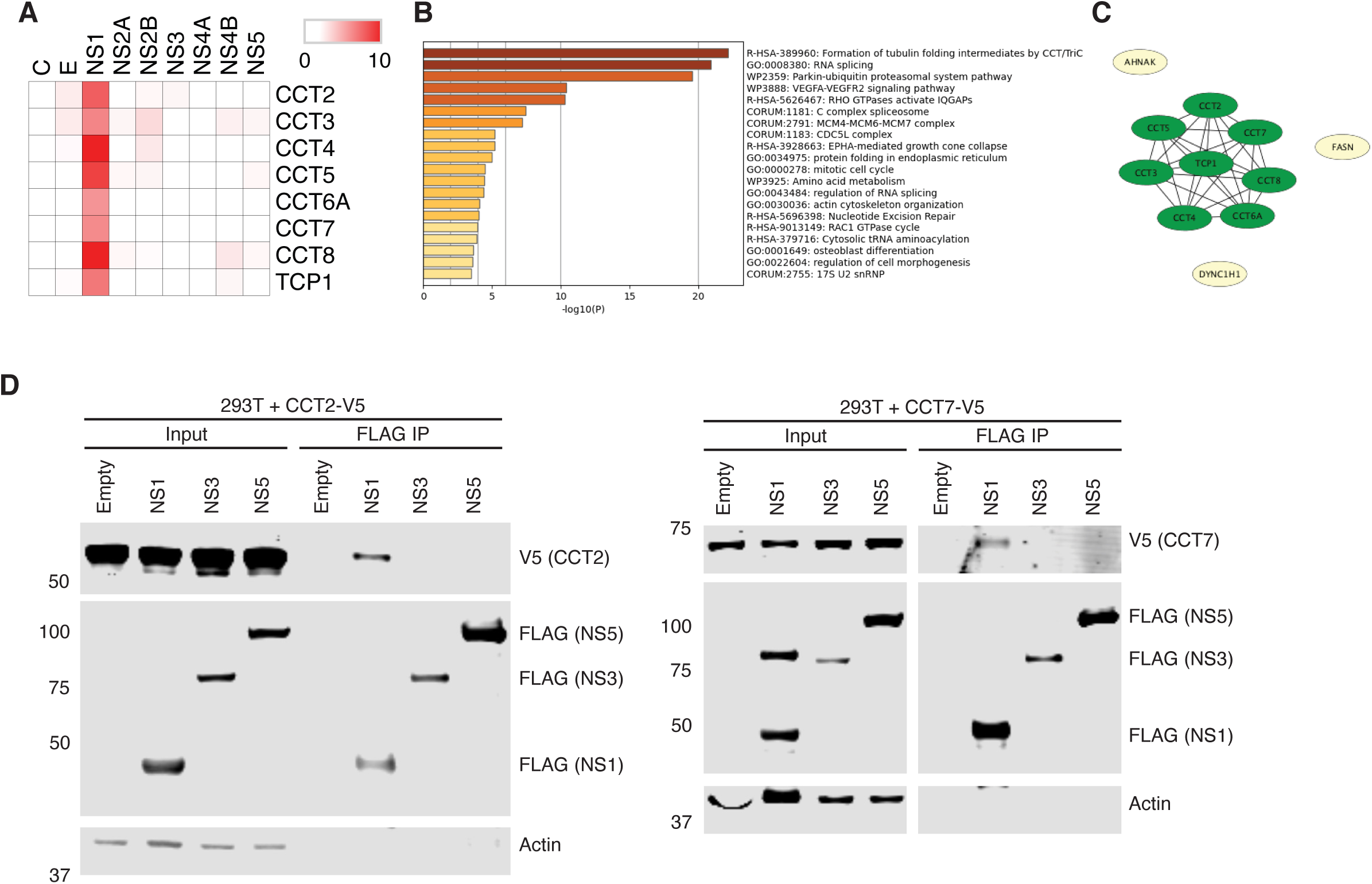
The CCT complex interacts specifically with DENV NS1. **a**. Heat map depicting the average CRAPome scores of the detected CCT complex members from the biological replicates of each bait DENV protein. **b.** Bar graph showing the enriched terms associated with all identified NS1 interacting proteins. Metascape analysis was performed as in Figure 2. **c.** Protein-protein interaction network showing the interactions between DENV NS1 and its PIPs. **d.** FLAG immunoprecipitation of 293T cells transfected with the indicated FLAG-tagged DENV proteins and V5-tagged CCT complex members, followed by Western blotting with the indicated antibodies, showing a specific interaction between the CCT complex and DENV NS1.

**Extended Data Fig. 4.**
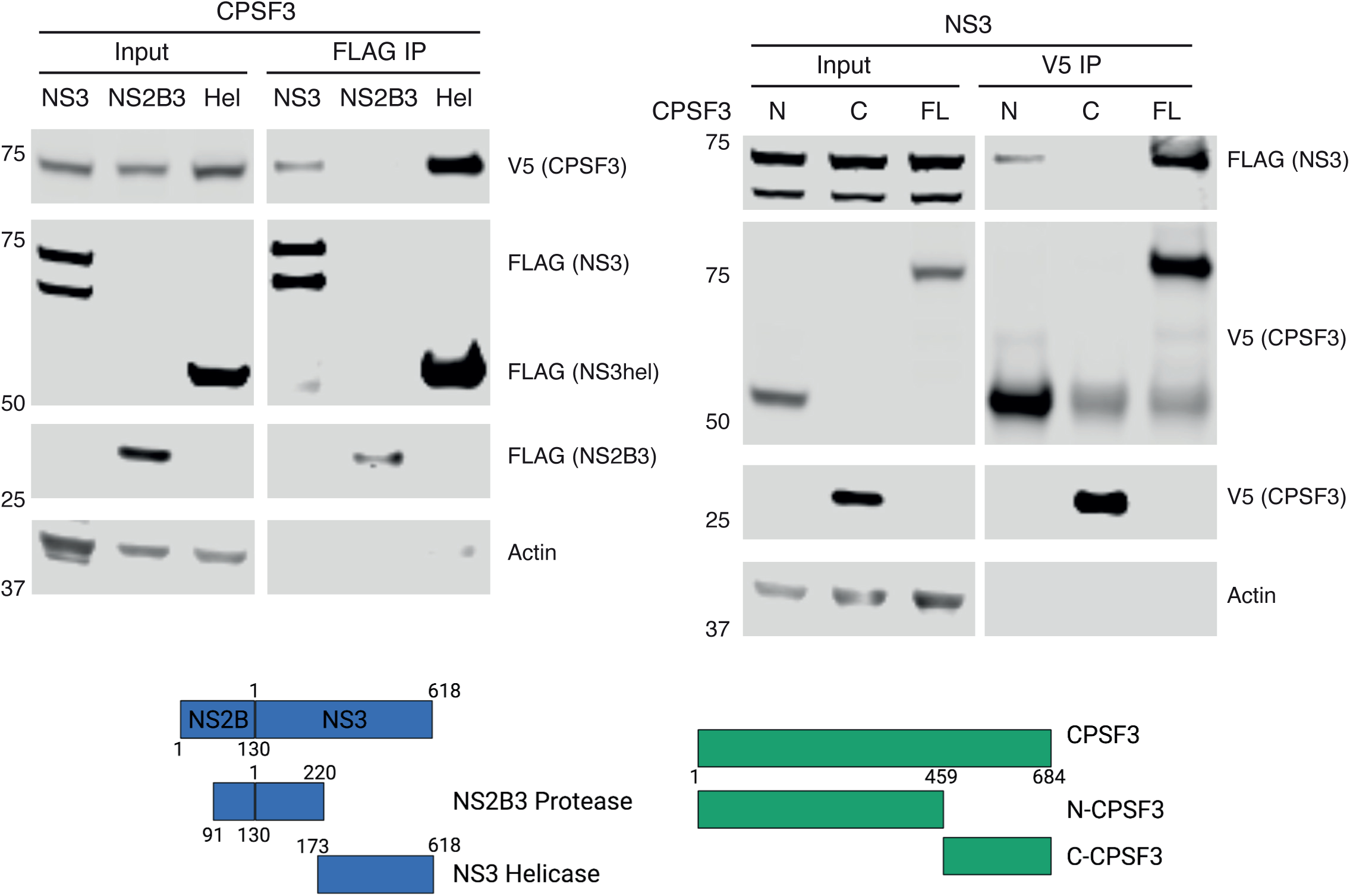
The helicase domain of DENV NS3 is necessary and sufficient to bind the N-terminal of CPSF3. Immunoprecipitation using the indicated antibodies of 293T cells transfected with FLAG-tagged NS3 and V5-tagged CPSF3, followed by Western blotting with the indicated antibodies. The amino acid positions for the various domains for DENV NS3 and CPSF3 used are depicted.

